# Nanoclustering and signaling of KRAS G12C and KRAS G12D respond to lipid acyl chain remodeling in an allele-specific manner

**DOI:** 10.1101/2024.05.30.596653

**Authors:** Neha Arora, Hong Liang, Walaa Kattan, Wantong Yao, Haoqiang Ying, Junchen Liu, Yong Zhou

## Abstract

Small GTPase KRAS mutated at hotspots, such as G12, G13 and Q61, are major drivers of cancer and display allele-specific oncogenic properties, which are not well understood. KRAS mutants require precise spatiotemporal distribution to the proteolipid nanoclusters on the plasma membrane (PM) for efficient signaling. We recently reported allele-specific lipid sensing of KRAS mutants. KRAS^G12D^, KRAS^G12V^ and KRAS^Q61H^ favor the unsaturated phosphatidylserine (PS), while KRAS^G12C^ and KRAS^G13D^ gain enrichment of the saturated PS, cholesterol and/or phosphoinositol 4,5-bisphosphate (PIP_2_). We, here, examined how the allele-specific lipid sensing of KRAS mutants contributes to their allele-specific signaling and activities. We now show that the stable expression of lysophosphatidylcholine acyltransferase 1 (LPCAT1) elevates the saturated phospholipids and reduces the mixed-chain lipids, especially PS species. Our super-resolution electron microscopy (EM)–spatial analysis revealed that the LPCAT1 expression perturbs the PM nanoclustering of KRAS^G12D^, without affecting that of KRAS^G12C^. LPCAT1 suppresses the KRAS-dependent mitogen-activated protein kinases (MAPKs) signaling and the MAPK-regulated proliferation and colony formation of the KRAS^G12D^-expressing human pancreatic PANC1 cells, while promoting those of the KRAS^G12C^-expressing MiaPaCa-2 cells. Mouse embryonic fibroblasts (MEF) transformed with KRAS^G12C^ contain more saturated lipids than those expressing KRAS^G12D^. Concordantly, patient tumor genomics analysis illustrated that expression of LPCAT1 and KRAS mutants negatively correlate in pancreatic adenocarcinoma with KRAS^G12D^ as a dominant driver, but loses correlation in lung adenocarcinoma with KRAS^G12C^ as a major driver. Thus, the allele-specific lipid sensing of KRAS mutants contributes to their pathological activities.

## Introduction

KRAS4B (or KRAS) small GTPase is a molecular switch that toggles between the inactive GDP-bound and active GTP-bound states (1–4). KRAS activates a wide variety of signaling cascades, especially mitogen-activated protein kinases (MAPKs) and phosphoinositol 3 kinase (PI3K), and regulates cell survival, growth, division, proliferation and migration (1–4). Constitutively active mutants of KRAS at hotspot residues G12, G13 and Q61 in its enzymatic globular G-domain (amino acids 1-166) are major drivers of cancer, contributing to 98% of pancreatic, 45% of colorectal and 31% of lung tumors (1–4). The discovery of allosteric pockets on the G-domain inspired the development of the allosteric inhibitors that effectively target specific KRAS mutants (5, 6). Interestingly, we recently reported that the G-domain allostery also contributes to the spatiotemporal organization of KRAS (7). A hallmark of KRAS pathophysiology is the compartmentalization of its oncogenic signaling to the proteolipid nanoclusters on the inner leaflet of the plasma membrane (PM) (8, 9). While its C-terminal membrane-anchoring domain has been mostly attributed to the membrane association of KRAS (9, 10), its peripheral G-domain only weakly touches the membranes and is generally overlooked for its roles in the spatiotemporal organization of KRAS. The G-domain allostery results in distinct G-domain / membrane interfaces (5, 11–17). We reported that key basic residues at the G-domain / membrane interfaces play distinct roles in the sensing of acyl chain structures of phosphatidylserine (PS) species (7). We further illustrated that KRAS oncogenic mutants, which undergo allele-specific allosteric reorientation (18–20), selectively associate with distinct PM lipids. KRAS^G12D^, KRAS^G12V^ and KRAS^Q61H^ possess remarkable selectivity for the mixed-chain phosphatidylserine (PS) species, while KRAS^G12C^ and KRAS^G13D^ gain association with the dual-saturated and dual-unsaturated PS species, cholesterol and/or phosphoinositol 4,5-bisphosphate (PIP_2_) (7). The distinct lipids enriched in these nanoclusters are functionally important because effectors of KRAS possess their own lipid-recognizing motifs and require synergistic binding of both KRAS and select lipids for efficient PM association and signal propagation (1, 2, 21). Thus, the allele-specific oncogenesis of KRAS mutants may depend on their allele-specific lipid sensing, as well as remodeling of lipidomes.

Here, we examined how remodeling homeostasis of the saturated/mixed-chain lipids may differentially impact activities of KRAS mutants. Lysophosphatidylcholine acyltransferases (LPCATs) remodel the *sn-2* acyl chains of phospholipids, with LPCAT1 preferentially catalyzing the generation of the dual-saturated lipids, especially dipalmitoylphosphatidylcholine (DPPC or di16:0 PC) (22, 23). We now show that the stable expression of LPCAT1 elevates the saturated phosphatidylcholine (PC) and phosphatidylethanolamine (PE), and depletes major mixed-chain PS species in baby hamster kidney (BHK) cells and human pancreatic ductal adenocarcinoma (PDAC) MiaPaCa-2 cells. The LPCAT1 expression more effectively disrupts the PM association, signaling and oncogenic activities of KRAS^G12D^, while not affecting or even promoting those of KRAS^G12C^. Taken together, oncogenic activities of KRAS mutants depends on lipid metabolism in an allele-specific manner.

## Results

### LPCAT1 elevates saturated phospholipids and depletes unsaturated phospholipids

We generated BHK cells and human PDAC MiaPaCa-2 cell lines stably expressing empty vector V2 or LPCAT1, which was validated in Western blotting (Fig.S1). The subsequent shotgun lipidomics illustrated that BHK cells expressing LPCAT1 contained significantly higher levels of the saturated PC and PE species (Fig.1 and B). Specifically, multiple saturated PC/PE species, including di14:0 PC, di16:0 PC, 14:0/18:0 PE and 16:0/18:0 PE, were particularly elevated in BHK cells stably expressing LPCAT1 (Fig.2A and B), consistent with previous studies (22, 24). Furthermore, the LPCAT1 expression markedly reduced the mixed-chain phospholipids with monounsaturated *sn-2* acyl chains, especially PS, phosphoinositols (PI), sphingomyelin (SM) and ceramides (Cer) (Fig.1). We then focused on PS species as KRAS spatial distribution is particularly sensitive to PS lipids. As shown in Fig.2C, various major mixed-chain PS species, such as 18:0/18:1 PS, 18:0/18:2 PS, 18:0/20:1 PS, 18:0/20:2 PS, 18:0/22:3 PS and 18:0/22:5 PS, were significantly lower in BHK cells expressing LPCAT1 than cells expressing the V2 vector control. Similarly, MiaPaCa-2 cells expressing LPCAT1 contained higher levels of the saturated PC and PE species, especially 14:0/16:0 PC, di16:0 PC, 14:0/16:0 PE and 14:0/18:0 PE (Fig.S2A and B). The LPCAT1 stable expression also depleted the mixed-chain PS species with monounsaturated and polyunsaturated *sn-2* chains, such as 18:0/18:1 PS (comprising 28% of the total PS), 18:0/18:2 PS (9% of the total PS) and 18:0/20:3 PS (28% of the total PS) (Fig.S2C). Taken together, the LPCAT1 stable expression elevates levels of the saturated phospholipids and depleted the mixed-chain PS species with monounsaturated and polyunsaturated *sn-2* chains.

**Figure 1.**
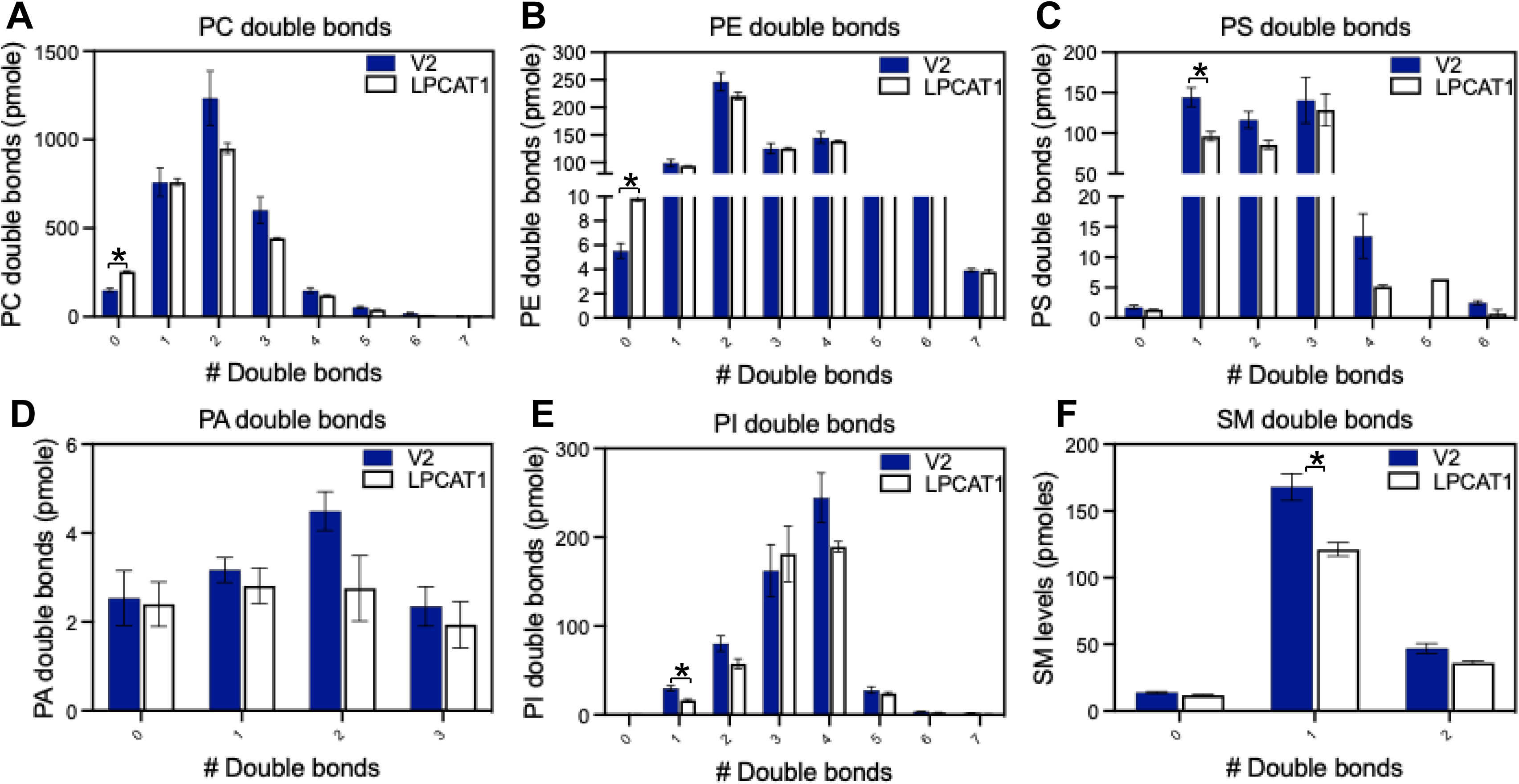
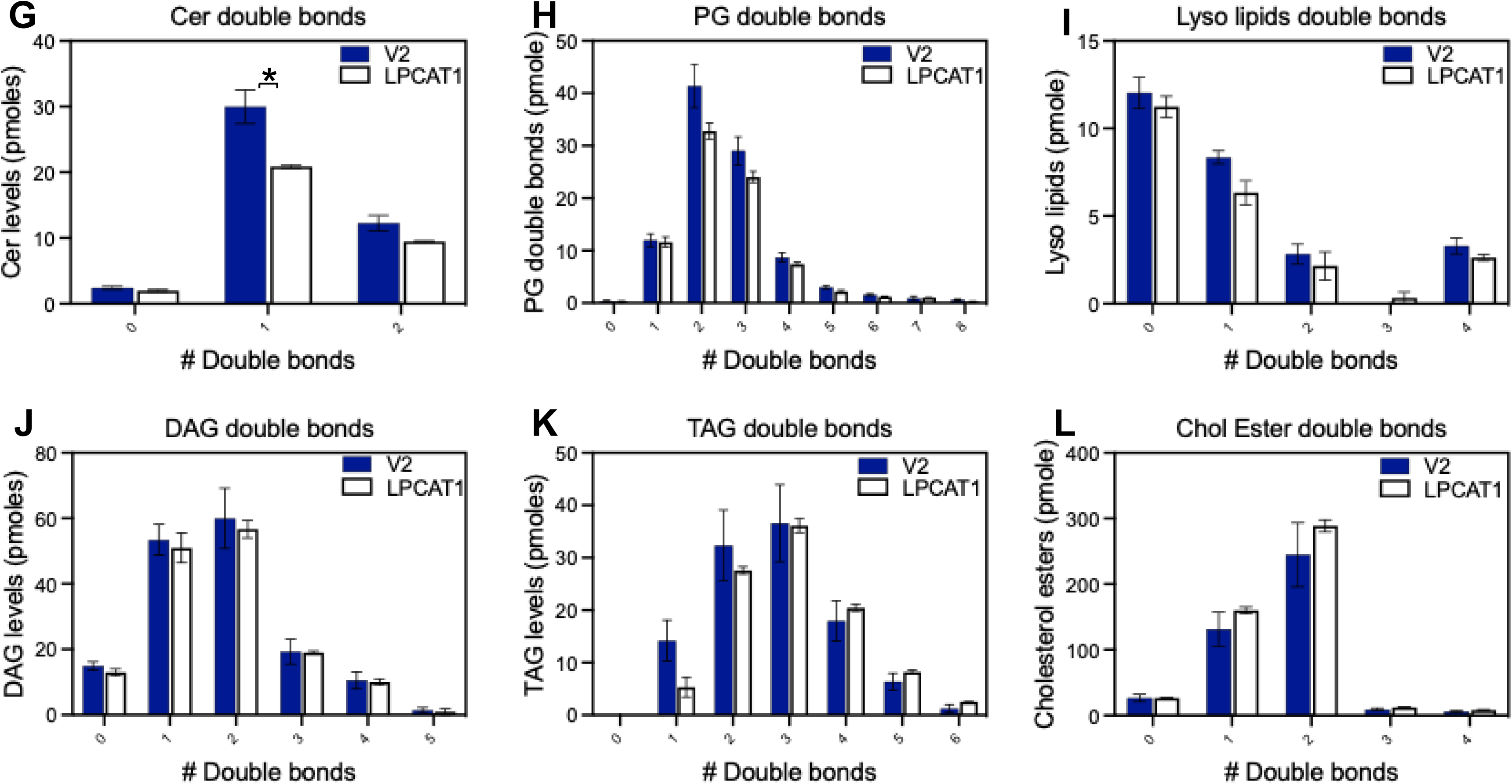
LPCAT1 expression elevates the saturated lipids in mammalian cells. Whole-cell lysates of baby hamster kidney (BHK) cells stably expressing V2 empty vector or Lpcat1 were collected for shotgun lipidomics. Levels of lipid species with different double bonds (in pmoles) for various lipid types are shown as mean ± SEM from 3 independent trials. Student’s t-test was used to evaluate the statistical significance, with * indicating p < 0.05.

**Figure 2.**
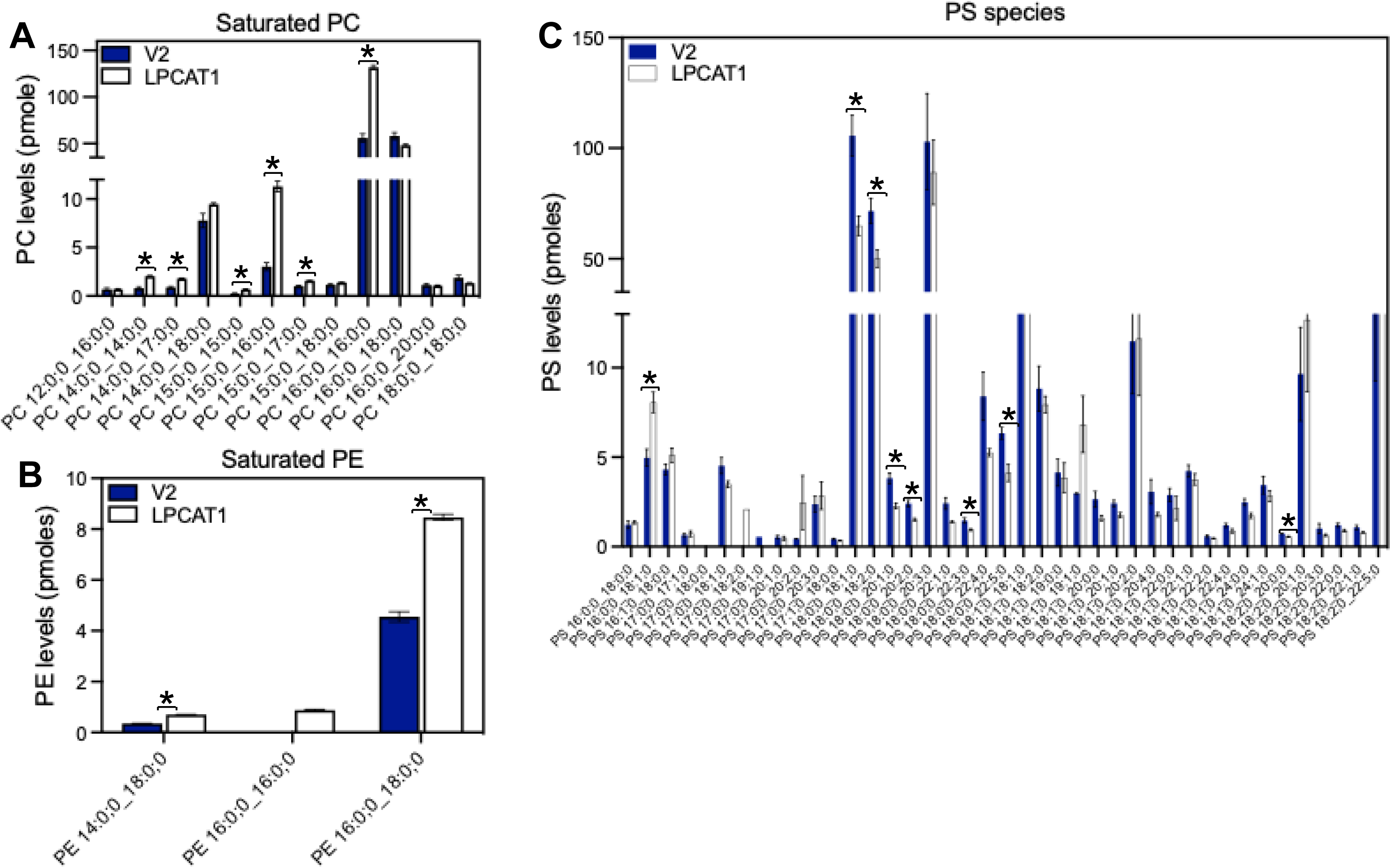
LPCAT1 alters homeostasis of main phospholipid types. Stable expression of V2 or Lpcat1 elevated the levels of multiple saturated PC (A) and PE (B) species. (C) Effects of V2 or Lpcat1 expression on the levels of all the PS species are shown. Data are shown as mean ± SEM from 3 independent trials. Student’s t-test was used to evaluate the statistical significance, with * indicating p < 0.05.

### LPCAT1 differentially impacts the PM association of KRAS mutants

We recently reported the allele-specific lipid sensing of KRAS oncogenic mutants, with KRAS^G12D^ and KRAS^G12V^ preferring the mixed-chain PS and KRAS^G12C^ gaining association with the dual-saturated and dual-unsaturated PS species, cholesterol and/or PIP_2_ (7). To test how KRAS mutants might respond to lipid acyl chain remodeling, we compared effects of the LPCAT1 stable expression on the signaling nanoclusters of GFP-KRAS^G12C^ and GFP-KRAS^G12D^, using electron microscopy (EM)-univariate nanoclustering analysis. Briefly, BHK cells stably expressing V2 or LPCAT1 were ectopically transfected with GFP-KRAS^G12C^ or GFP-KRAS^G12D^. The apical PM sheets of these cells were attached to copper EM grids. GFP-KRAS mutants anchored to the PM inner leaflet were immunolabeled with anti-GFP antibody conjugated to 4.5 nm gold nanoparticles. The gold-labeled GFP-KRAS mutants were imaged via transmission EM (TEM) at 100,000x magnification. Spatial distribution of gold particles within a 1 μm^2^ PM area was quantified using the Ripley’s K-function analysis, where the extent of nanoclustering, *L*(*r*) – *r*, was plotted against distance *r* in nanometers. The peak *L*(*r*) – *r* value, or *L_max_*, was used as a statistical summary for the nanoclustering. *L*(*r*) – *r* values above the 99% confidence interval (99% CI) of 1 indicate the statistically meaningful nanoclustering, with the larger *L_max_* values corresponding to more extensive nanoclustering. The number of gold particles within the same1 μm^2^ PM area estimates the PM localization of GFP-KRAS. In Fig.3A, *L_max_* of GFP-KRAS^G12C^ was similar in BHK cells expressing either V2 or LPCAT1, indicating that the LPCAT1 expression had minimal effect on the nanoclustering of KRAS^G12C^. The nanoclustering (*L_max_*) of GFP-KRAS^G12D^ was markedly lower in BHK cells expressing LPCAT1 than the V2 vector control, indicating that the nanoclustering of KRAS^G12D^ is more sensitive to the perturbation of LPCAT1 expression. In Fig.4B, the gold labeling of GFP-KRAS^G12C^ decreased by ∼ 35% in LPCAT1-expressing cells when compared to the V2 control, while the gold labeling of GFP-KRAS^G12D^ decreased by ∼69% by LPCAT1 expression. Taken together, the PM association of KRAS^G12D^ is more sensitive to the LPCAT1 perturbation than that of KRAS^G12C^ and KRAS^G12D^.

**Figure 3.**
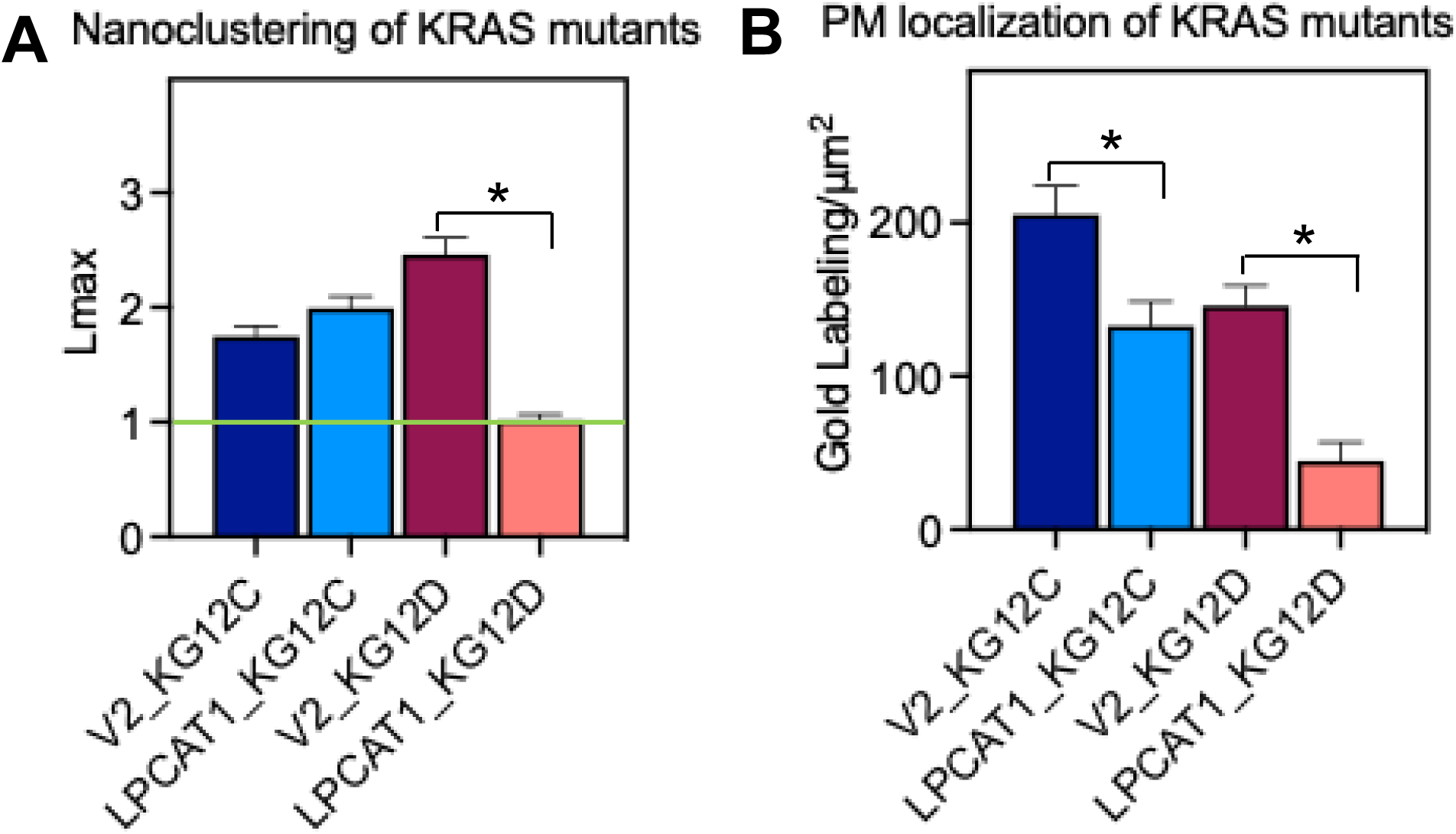
LPCAT1 preferentially disrupts the signaling nanoclustering of KRAS^G12D^ than KRAS^G12C^. Spatial distribution of KRAS mutants was quantified via electron microscopy (EM)-spatial analysis. Intact apical PM sheets of BHK cells stably expressing V2 or LPCAT1 transiently expressing GFP-KRAS^G12C^ or GFP-KRAS^G12D^ were attached to copper EM grids. GFP anchored to the PM inner leaflet was immunolabeled with anti-GFP antibody conjugated to 4.5 nm gold nanoparticles. Distribution of the gold-labeled GFP-KRAS^G12C^ and GFP-KRAS^G12D^ within a selected 1μm^2^ PM area was calculated using the Ripley’s K-function analysis. A nanoclustering curve was plotted as the extent of nanoclustering, *L*(*r*) – *r*, vs. length scale, *r* in nanometers. The peak value of the curve, termed as *L_max_*, was used as a summary statistic to indicate nanoclustering (A). The *L*(*r*) – *r* of 1 is the 99% confidence interval (99% CI, green line), the values above which indicate statistically meaningful clustering. Number of gold particles within the 1μm^2^ PM area was counted to indicate PM localization (B). The nanoclustering and PM localization of GFP-KRAS^G12C^ and GFP-HRAS^G12D^ of BHK cells are shown as mean ± SEM. For the nanoclustering data, the statistical significance was evaluated via the non-parametric bootstrap tests. For the gold labeling data, the statistical significance was quantified using the one-way ANOVA. * indicates p < 0.05.

**Figure 4.**
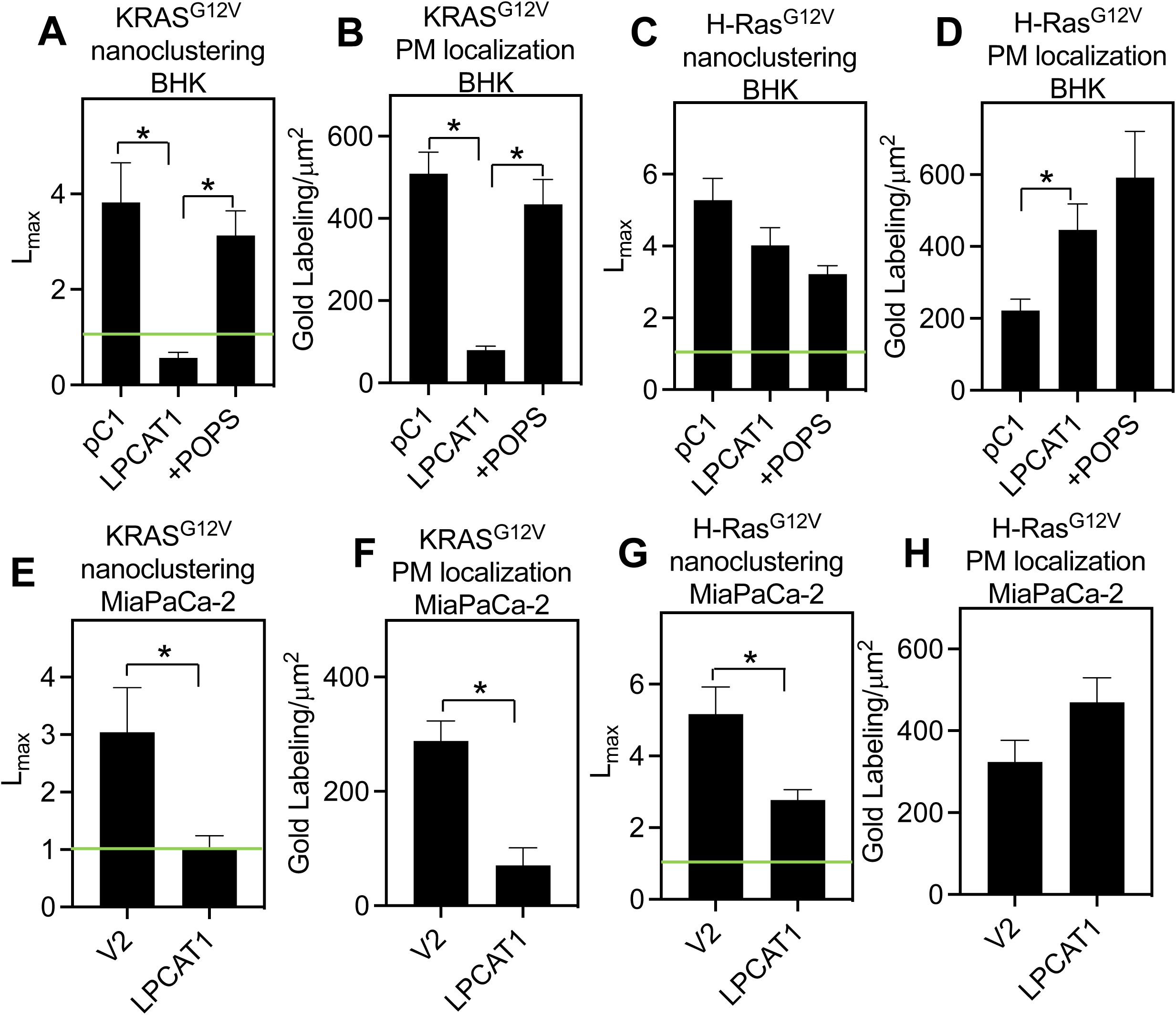
LPCAT1 more preferentially disrupts the signaling nanoclusters of KRAS^G12V^ on the plasma membrane. Spatial distribution of mammalian cells, including human pancreatic ductal adenocarcinoma MiaPaCa-2 and BHK cells, were quantified via EM-spatial analysis. Intact PM sheets of BHK (A-D) and MiaPaCa-2 (E-H) cells stably expressing V2 or LPCAT1 transiently expressing GFP-KRAS^G12V^ or GFP-HRAS^G12V^ were attached to EM grids. GFP anchored to the PM inner leaflet was immunolabeled with anti-GFP antibody conjugated to 4.5 nm gold nanoparticles. Distribution of the gold-labeled GFP-KRAS^G12V^ and GFP-HRAS^G12V^ within a selected 1μm^2^ PM area was calculated using the Ripley’s K-function analysis. The summary statistic for nanoclustering *L_max_* (A, C, E and G) and gold labeling density (B, D, F and H) are shown as mean ± SEM. For the nanoclustering data, the statistical significance was evaluated via the non-parametric bootstrap tests. For the gold labeling data, the statistical significance was quantified using the one-way ANOVA. * indicates p < 0.05.

We previously showed that KRAS^G12V^ and KRAS^G12D^ favor similar mixed-chain PS, while HRAS^G12V^ and KRAS^G12C^ gain association with all PS species tested (7). We next compared effects of LPCAT1 on the PM association of GFP-KRAS^G12V^ and GFP-HRAS^G12V^ in BHK cells. As expected, the LPCAT1 expression significantly disrupted the nanoclustering and PM localization of GFP-KRAS^G12V^, which was effectively restored by the acute addback of the mixed-chain 16:0/18:1 PS (POPS, Fig.4A and B). LPCAT1 did not impact the nanoclustering of GFP-HRAS^G12V^, while increasing the PM localization of GFP-HRAS^G12V^ (Fig.4C and D). Acute addback of POPS did not restore the PM association of GFP-HRAS^G12V^ in BHK cells. We observed similarly differential effects of LPCAT1 on the PM association of GFP-tagged KRAS^G12V^ and HRAS^G12V^ in human PDAC MiaPaCa-2 cells (Fig.4E-H). Taken together, the LPCAT1-induced lipid acyl chain remodeling differentially impacts the PM association of KRAS mutants in distinct manners. *LPCAT1 impacts MAPK signaling and oncogenic activities of KRAS-dependent tumor cells in an allele-specific manner*.

We next compared effects of LPCAT1 on signal output of MAPK and PI3K cascades in the KRAS-dependent human PDAC cell lines expressing different KRAS mutants, including MiaPaCa-2 (KRAS^G12C^), MOH (KRAS^G12R^) and PANC1 (KRAS^G12D^) cells, and the wild-type KRAS-expressing BxPC3 cells. The stable LPCAT1 expression resulted in a ∼ 50% increase in the level of the phosphorylated ERK (pERK/total ERK) in MiaPaCa-2 cells, while significantly decreased pERK/total ERK levels in MOH and PANC1 cells and having minimal effect on MAPK signaling in BxPC3 cells (Fig.5A-E). The PI3K signaling (pAkt/total Akt) less preferentially regulated by KRAS was unaffected by LPCAT1 expression (Fig.5A, F-I).

**Figure 5.**
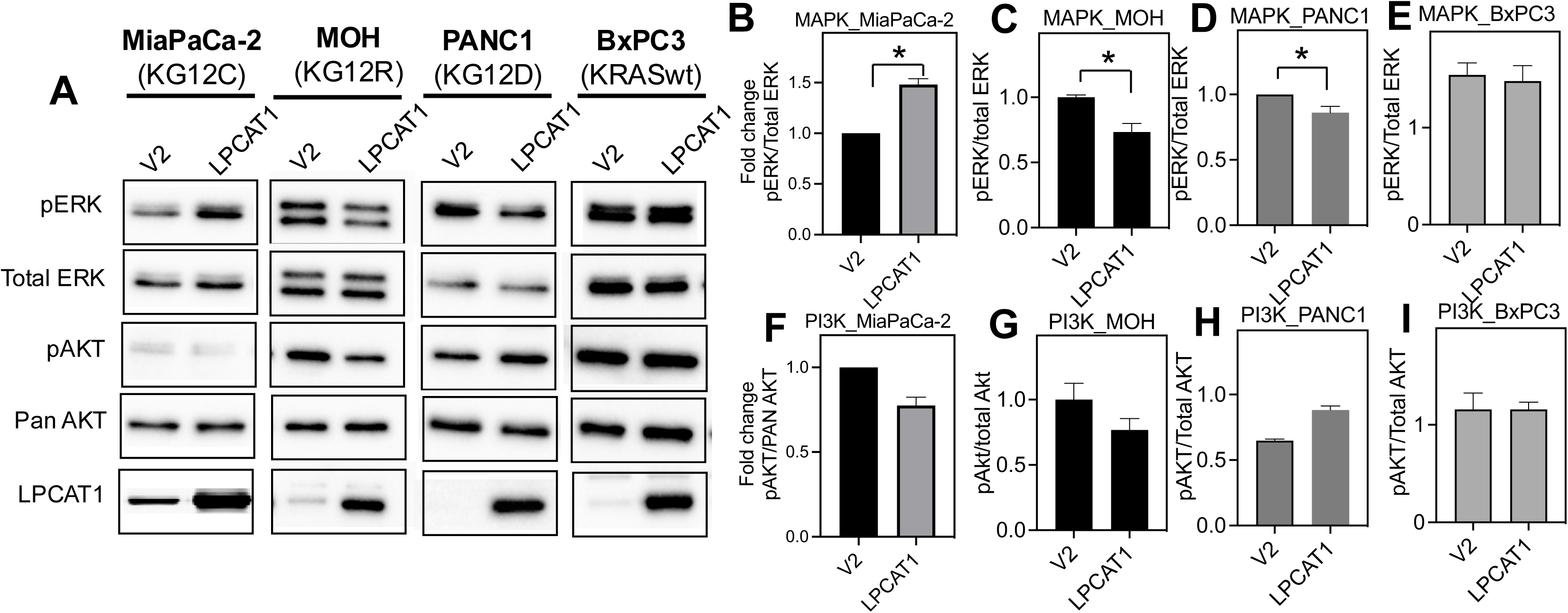
LPCAT1 differentially impacts the signaling and oncogenic activities of KRAS mutants in an allele-specific manner. (A) Whole-cell lysates of human pancreatic tumor lines, including MiaPaCa-2 (KRAS^G12C^), MOH (KRAS^G12R^), PANC1 (KRAS^G12D^) and BxPC3 (KRAS wild-type) stably expressing V2 or LPCAT1, were collected for Western blotting. Antibodies against the phosphorylated ERK (pERK), total ERK, pAkt, total Akt and LPCAT1 were used to blot for targeted proteins. Sample blots for a single trial are shown. Quantifications of pERK/total ERK for MiaPaCa-2 (B), MOH (C), PANC1 (D) and BxPC3 (E), as well as pAkt/total Akt for MiaPaCa-2 (F), MOH (G), PANC1 (H) and BxPC3 (I), are shown as mean ± SEM pooled from 3 independent experiments. Statistical significance was evaluated using Student’s t-test, with * indicating p < 0.05.

We then compared effects of LPCAT1 on oncogenic activities of MiaPaCa-2, MOH and BxPC3 cells. When compared with the V2 vector controls, the LPCAT1 stable expression increased the number of colonies of MiaPaCa-2 cells, while significantly reduced the colony formation of MOH cells and having no effect on colony formation of BxPC3 cells (Fig.6A-C). Similarly, the LPCAT1 stable expression increased the proliferation of MiaPaCa-2 cells, decreasing proliferation of MOH cells, while having no effect on BxPC3 cells (Fig.6D-F). Taken together, LPCAT1 differentially impacts signaling and oncogenic activities of the KRAS-dependent tumor lines in an allele-specific manner, consistent with its allele-specific effects on the spatiotemporal organization of KRAS mutants.

**Figure 6.**
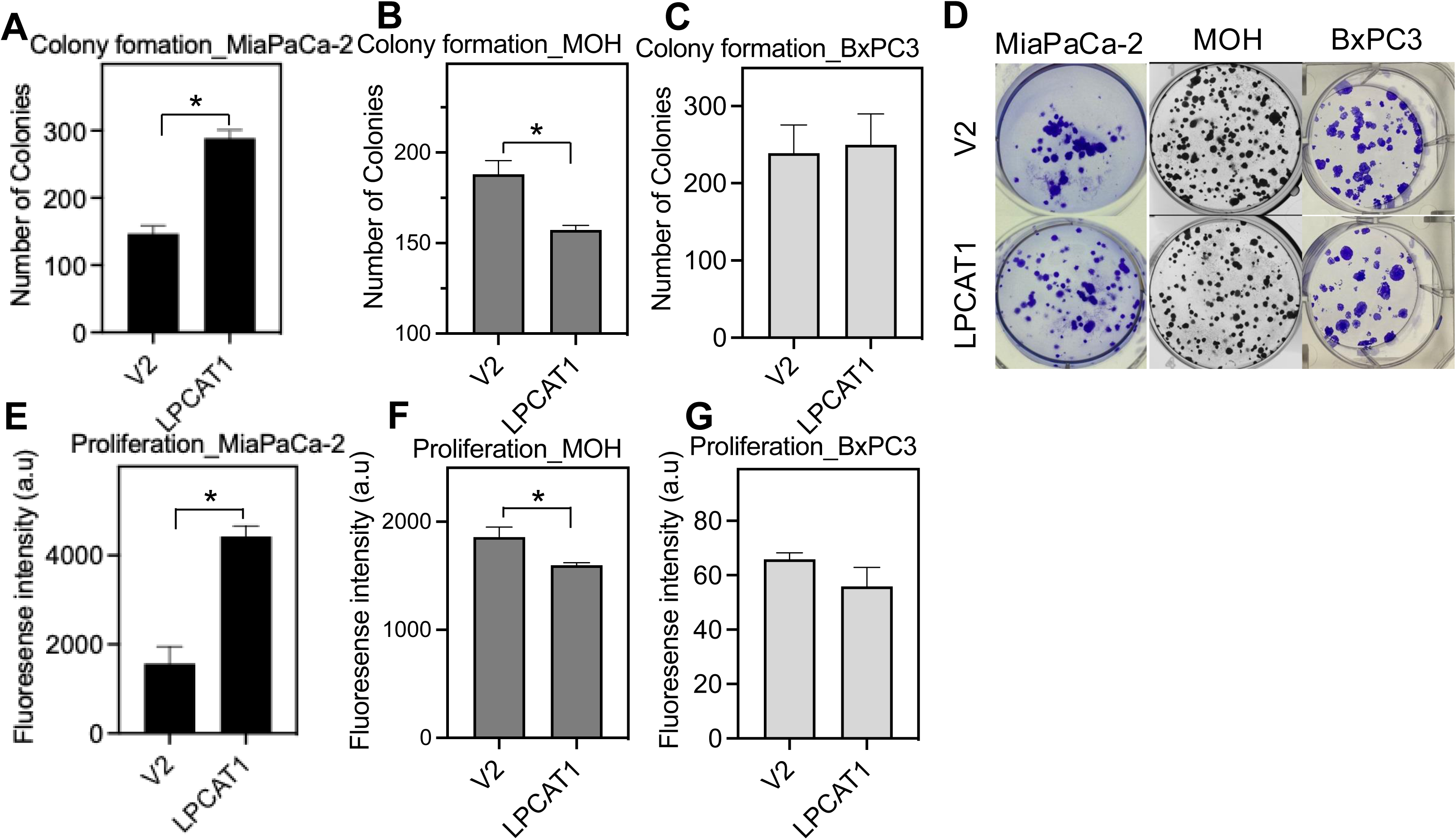
LPCAT1 differentially impacts oncogenic activities of tumor cells driven by different KRAS mutants. The KRAS-dependent and -independent human PDAC cells stably expressing V2 or LPCAT1 were seeded in 6-well plates. Colonies were counted after 96 hours of growth. The number of colonies for MiaPaCa-2 (A), MOH (B) and BxPC3 (C) are shown as mean ± SEM pooled from 3 independent trials. (D) Sample images of MiaPaCa-2, MOH and BxPC3 colonies are shown. To evaluate proliferation, MiaPaCa-2 (E), MOH (F) and BxPC3 (G) cells stably expressing V2 or LPCAT1 were seeded in 96-well plates. After 96 hours of growth, CyQUANT cell proliferation assay was used to measure proliferation. Data are shown as mean ± SEM pooled from 3 independent experiments. Data are shown as mean ± SEM pooled from 3 independent trials. For all experiments, Student’s t-test was used to evaluate the statistical significance with * indicating p < 0.05.

### KRAS^G12C^- and KRAS^G12D^-expressing cells possess distinct lipidomes

As seen above, the nanoclustering, signaling and oncogenic activities of KRAS mutants respond to lipid acyl chain remodeling in an allele-specific manner. It is possible that cells expressing different KRAS mutants might modulate their lipid profiles to more efficiently facilitate the activities of specific KRAS mutants. To test this, we compared lipidomes of RASless mouse embryonic fibroblasts (RASless MEF) expressing KRAS^G12C^ or KRAS^G12D^. The RASless MEFs are isolated from mice with null NRAS and HRAS alleles and a floxed KRAS locus, then infected with Cre Recombinase Adenovirus carrying genes of specific KRAS mutants (25). With all endogenous RAS genes deleted and distinct RAS mutants expressed, RASless MEF lines are ideal tools for comparing pathophysiological activities of specific RAS mutants (25). The whole-cell lysates of the RASless MEF lines expressing KRAS^G12C^ or KRAS^G12D^ were collected for lipidomics analysis. In Fig.7A-D, the RASless MEF cells expressing KRAS^G12C^ contained higher levels of the saturated PE and PI species than the KRAS^G12D^-expressing RASless MEF line, without affecting the total PC and PE levels. For PS species, despite similar total PS levels (Fig.7F), the KRAS^G12C^-expressing MEF cells contained distinct profiles of PS species than the KRAS^G12D^-expressing cells (Fig.7E). KRAS^G12D^ cells contained higher levels of the mixed-chain PS species with the fully saturated *sn-1* chain and polyunsaturated *sn-2* chains, such as 18:0/20:4 PS, 18:0/21:3 PS and 18:0/22:3 PS than KRAS^G12C^ cells. On the other hand, KRAS^G12C^ cells contained more mixed populations of different types of PS species including both the dually unsaturated PS species (18:1/18:3 PS, 18:1/20:3 PS and 18:1/24:1 PS) and the mixed-chain PS species with a saturated *sn-1* chain (16:0/17:1 PS, 16:0/18:3 PS, 18:0/18:3 PS). This is consistent with our previous findings that KRAS^G12D^ more selectively associates with the mixed-chain PS with a saturated *sn-1* chain, while KRAS^G12C^ associates with all PS species tested including both the mixed-chain and symmetric PS species (7). Taken together, the KRAS^G12D^-expressing cells more selectively enrich the mixed-chain PS species, while KRAS^G12C^-expressing cells partially lose such selectivity.

**Figure 7.**
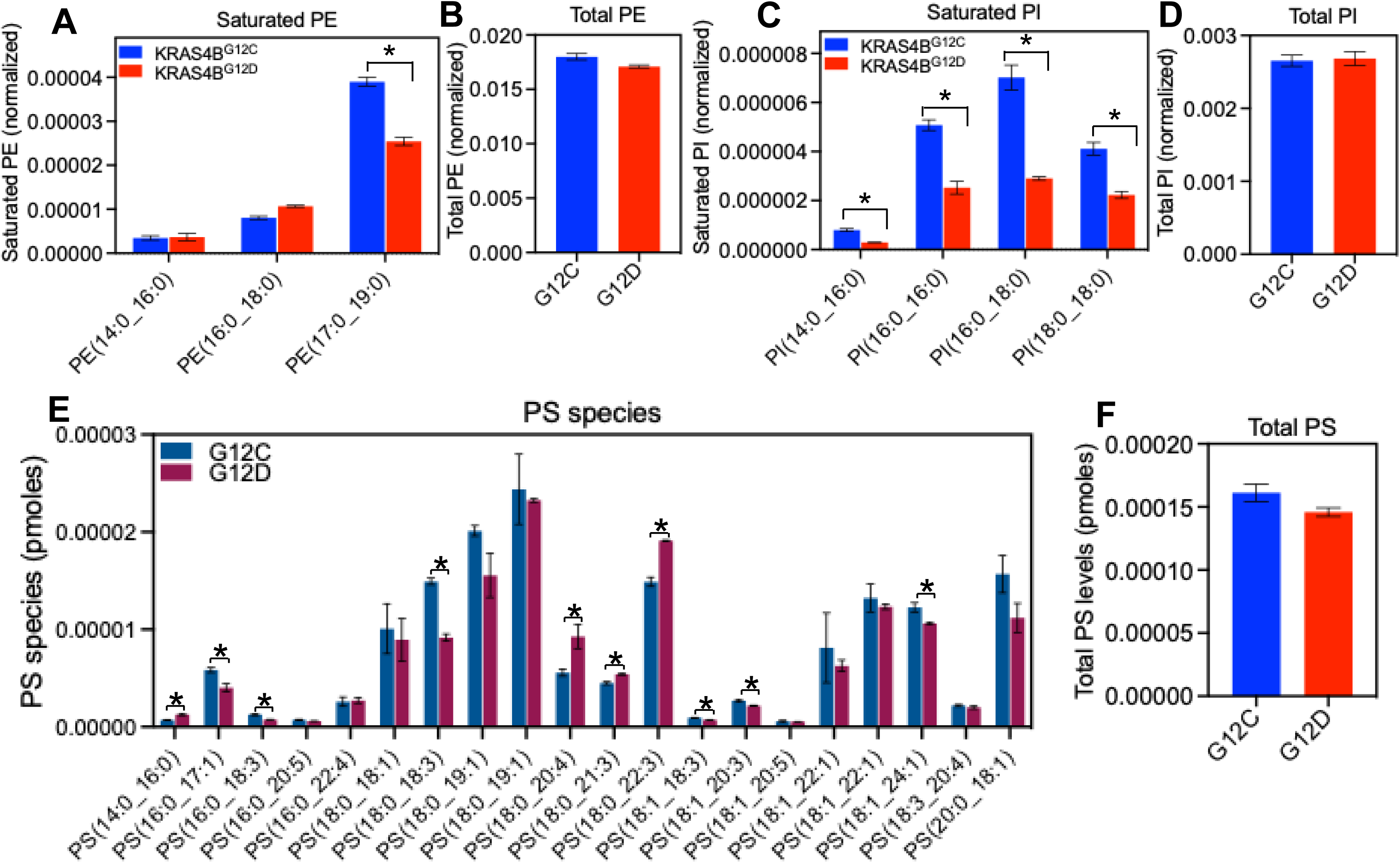
RASless mouse embryonic fibroblasts (MEFs) expressing KRAS^G12C^ contain more saturated lipids than RASless MEFs containing KRAS^G12D^. Whole-cell lysates of RASless MEFs driven by KRAS^G12C^ or KRAS^G12D^ were collected for lipidomics. Normalized levels of saturated PE (A), total PE (B), saturated PI (C) and total PI (D) are shown. Individual species of PS lipids (E) and total PS (F) are also shown. All data are shown as mean ± SEM pooled from 3 independent experiments. Student’s t-test was used to evaluate the statistical significance with * indicating p < 0.05.

### LPCAT1 expression negatively correlates with KRAS mutations in pancreatic adenocarcinoma (PAAD) but not lung adenocarcinoma (LUAD)

KRAS^G12D^ (46%) and KRAS^G12V^ (31%) are dominant mutations, while KRAS^G12C^ is a minor (3%), in human pancreatic adenocarcinoma (PAAD). On the other hand, KRAS^G12C^ (37%), KRAS^G12D^ (19%) and KRAS^G12V^ (22%) are equivalently prevalent in lung adenocarcinoma (LUAD). Thus, LPCAT1 may be more correlated with KRAS mutant expression in PAAD than LUAD. To further examine this, we compared The Cancer Genomic Atlas (TCGA) at the Genomic Data Commons (GDC). Fig.8A shows that LPCAT1 mRNA levels were significantly higher in PAAD patients with KRAS wild-type than those with KRAS mutations, suggesting that LPCAT1 expression negatively correlates with KRAS oncogenesis in PAAD. Interestingly, LPCAT1 expression did not correlate with KRAS mutations in LUAD patients (Fig.8B).

**Figure 8.**
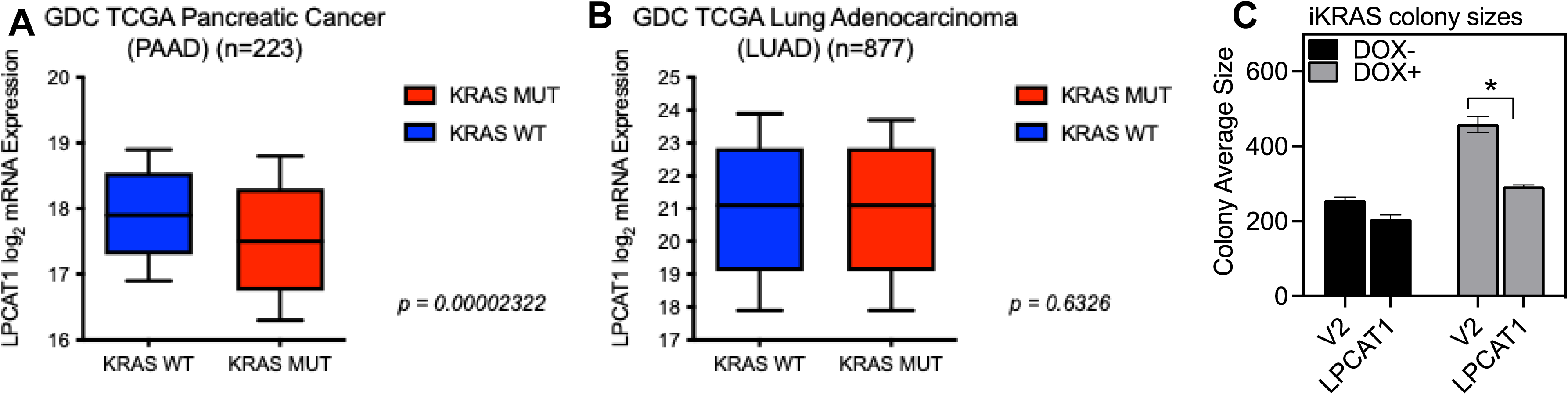
LPCAT1 expression correlates with KRAS mutant expression in pancreatic adenocarcinoma but not lung adenocarcinoma. Statistical analyses were performed using patient data obtained from the Cancer Genomic Atlas (TCGA) in the Genomic Data Commons (GDC) data portal. LPCAT1 mRNA levels were compared in pancreatic adenocarcinoma (A) or lung adenocarcinoma (B) patients with the KRAS-dependent and -independent tumors. Welch’s t-tests were performed to evaluate the statistical significance. (C) To further validate the KRAS specificity of LPCAT1 expression, we used murine pancreatic adenocarcinoma iKRAS cells with inducible expression of KRAS^G12D^. iKRAS cells were maintained in doxycycline (DOX+) to induce expression of KRAS^G12D^, or withdrawn from DOX for 48 hours (DOX-) for KRAS independent condition. iKRAS (DOX+/-) cells were seeded in 6-well plates. After 96 hours, sizes of the colonies were measured. Data are shown as mean ± SEM pooled from 3 independent experiments. Student’s t-test was used to evaluate the statistical significance with * indicating p < 0.05.

To further evaluate the specificity of LPCAT1 in impacting KRAS mutants, we stably expressed V2 empty vector or LPCAT1 in an inducible KRAS^G12D^-driven PDAC iKRAS mouse line (26). Fig.8C illustrates that the LPCAT1 stable expression significantly decreased the sizes of colonies of iKRAS lines induced to express KRASG12D, while having minimal effects of iKRAS cells without KRAS mutant expression.

## Discussion

Allele specific oncogenesis of KRAS have been consistently observed. For instance, in pancreatic tumors, G12D and G12V mutations are highly prevalent, comprising ∼77% of all KRAS-driven pancreatic tumors, while G12C is a minor mutation accounting for 3% of the KRAS-dependent tumor formation in the pancreas (4, 27). In lung adenocarcinoma, G12C becomes markedly more dominant, contributing to 37% of tumor formation in the lung (4, 27). *In vitro* binding assays further demonstrated that KRAS mutants recruit effectors and undergo the GTPase activating protein (GAP)-mediated GTP hydrolysis in an allele-specific manner (28). The mechanisms mediating these allele-specific pathophysiological activities of KRAS mutants are not well-understood. We recently reported that KRAS oncogenic mutants recognize distinct lipids in an allele-specific manner (7). Despite being in the prolonged activated states, KRAS mutants must be localized to the proteolipid nanoclusters on the PM to efficiently propagate signaling. This is because most effectors of KRAS possess their own lipid-binding motifs and require synergistic association with both the activated KRAS and select lipids for proper signal transmission. In the current study, we compared how the allele-specific lipid sensing of KRAS^G12C^ and KRAS^G12D^ contributes to their distinct signaling and oncogenic activities. We now show that remodeling lipidomes of mammalian cells via stable expression of LPCAT1 (prefers the fully saturated fatty acid chains as substrates) differentially modulates the spatial distribution, signaling and oncogenesis of KRAS^G12C^ and KRAS^G12D^ in distinct manners.

While RAS isoforms are peripheral membrane proteins with their G-domains primarily localized to the outside of the membranes, their C-terminal membrane-anchoring domains have been mainly attributed as their membrane-associating features. As all constitutively active mutants of KRAS share identical membrane anchors, it has been traditionally considered that KRAS mutants prefer similar lipids and embed in similar nanoclusters in the non-rafty regions enriching the unsaturated lipids in the absence of cholesterol. However, it has long been observed that RAS proteins with the same membrane anchors can distribute to separate regions of the membranes. For instance, the GTP-and GDP-bound RAS isoforms are spatially segregated. For HRAS and NRAS, GTP-loading moves the isoforms between the cholesterol- and saturated lipid-enriched rafts and the highly fluid non-raft domains (8, 29). Although the GTP- ad GDP-bound KRAS both prefer the highly fluid non-rafty domains, they still segregate from each other (8). Abankwa et al. (30, 31) first proposed that HRAS G-domain can swing its G-domain towards or away from the membranes upon GTP-/GDP-exchange, which contributes its raft/non-raft shifts.

We recently reported that KRAS oncogenic mutants, most of which are prolonged GTP-bound, also prefer distinct lipids with headgroup and acyl chain specificity (7). Most surprisingly, KRAS^G12C^ and KRAS^G13D^ strongly favor the fully saturated PS, cholesterol and/or PIP_2_ in the PM. In accordance with these findings, we now illustrate that the stable expression of LPCAT1, with the saturated fatty acids as its favored substrates, differentially regulates the nanoclustering and function of KRAS^G12C^ and KRAS^G12D^. Although the G-domains of KRAS mutants mostly suspend outside of the membranes, we, here, present exciting evidence that the G-domain-mediated lipid sensing directly participates in signaling and oncogenic activities of KRAS.

Targeting lipid metabolism has been perceived as lacking specificity, thus potentially elevated cytotoxicity. Further, earlier attempts to inhibit the prenylation of KRAS via farnesyltransferase inhibitors (FTIs) have not been successful since KRAS mutants are alternatively geranylgeranylated in the presence of FTIs (32–34). This experience has considerably dampened the enthusiasm of targeting lipid metabolism as a potential strategy for inhibiting KRAS oncogenesis. We now show that the LPCAT1-mediated remodeling lipidomes via LPCAT1 selectively and differentially impacts different KRAS mutants. LPCAT1 expression has been shown to promote oncogenic activities of epidermal growth factor receptor (EGFR)-driven cancer by elevating levels of the fully saturated PC and promoting lipid rafts (23). This reflects the complex biological and pathological roles of LPCAT1 and intricate selectivity of lipid acyl chain remodeling. EGFR dimerization/oligomerization occur in lipid rafts enriched with cholesterol and saturated lipids, which in turn promotes autophosphorylation and signaling (23, 35). KRAS^G12C^ distributes to similar raft domains (7), thus its signaling and oncogenic activities are promoted upon LPCAT1 expression. On the other hand, KRAS^G12D^ and KRAS^G12V^ favor the mixed-chain PS species with unsaturated *sn-2* chains (7). Consistently, we now show that the LPCAT1 expression attenuates the nanoclustering and signaling of KRAS^G12D^ and KRAS^G12V^. Thus, the opposing effects of LPCAT1 on the cholesterol-dependent EGFR and KRAS^G12C^ vs. the cholesterol-poor KRAS^G12D^ and KRAS^G12V^ suggest potential selectivity, with which target lipid metabolism can achieve. Taken together, remodeling lipid acyl chains impacts cell signaling events on membranes in distinct manners.

## Conclusion

KRAS oncogenic mutants selectively sense distinct lipids. Altering homeostasis of PS species may be a novel alternative strategy to inhibit KRAS oncogenesis. Here, we show that higher LPCAT1 expression elevates the saturated lipids and depletes major unsaturated PS species, which differentially impacts the nanoclustering and signaling of KRAS oncogenic mutants. Thus, LPCAT1 may serve as a marker when considering treatment options for the KRAS-dependent cancer. In the future, specific promoters of LPCAT1, or inhibitors of LPCAT1 antagonists, may be explored as alternative treatment strategies for KRAS cancer.

## Acknowledgements

This work was supported in part by the National Institutes of Health R01GM138668 to N. Arora, H. Liang and Y. Zhou.

## Materials and Methods

### Electron microscopy (EM)-spatial analysis

#### EM-univariate nanoclustering

Apical or basolateral PM of baby hamster kidney (BHK) or human pancreatic tumor MiaPaCa-2 cells expressing GFP-KRAS^G12V^ or GFP-HRAS^G12V^ was attached to EM grids. The intact native PM sheets were then fixed with 4% paraformaldehyde (PFA) / 0.1% gluaraldehyde, tagged with anti-GFP antibody conjugated with 4.5 nm gold nanoparticles, and negative stained with 0.3% uranyl acetate, and embedded in methyl cellulose. Transmission EM (TEM) was used to image PM sheets at 100,000x magnification. ImageJ was then used to assign the x / y coordinates of each gold particle within a select 1μm^2^ PM area. Ripley’s K-function calculated the nanoclustering of the gold-labeled GFP-RAS on the PM. The null hypothesis of this analysis is that the gold nanoparticles distribute in a random pattern:

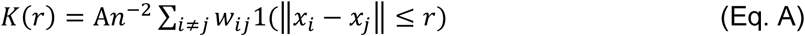

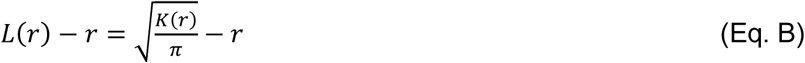

In Equation A, *K(r*) denotes the univariate distribution for gold nanoparticles with a total number of *n* in a PM area of *A*; *r* signifies the distance between gold particles with an increment of 1 nm from 1 to 240 nm; || · || denotes Euclidean distance that describes an indicator of 1( · ) = 1 if ||*x_i_*-*x_j_*|| ≤ r and 1( · ) = 0 if ||*x_i_*-*x_j_*|| > r. *w* ^-1^ is used to correct edge effects by describing the fraction of the circumference of a circle with the center defined as *x_i_* and radius ||*x_i_*-*x_j_*||. In Equation B, *L*(*r*) – *r* denotes the linear transformation of *K*(*r*) in Eq. A, which is achieved by normalizing *K*(*r*) against the 99% confidence interval (99% C.I.) calculated via Monte Carlo simulations. *L*(*r*) - *r* = 0 when gold nanoparticles distribute a complete random pattern. *L*(*r*) - *r* values above the 99% confidence interval (99% CI) of 1 indicate statistically meaningful clustering, with larger *L*(*r*) - *r* values describing more extensive clustering. The peak values of *L*(*r*) - *r* curves, termed as *L_max_*, are used as a summary statistic to signify the extent of nanoclustering. For each condition, at least 15 PM sheets from individual cells were imaged, analyzed and pooled. Statistical significance was evaluated via comparing our calculated point patterns against 1000 bootstrap samples in bootstrap tests (36, 37).

#### EM-Bivariate co-clustering analysis

The K-function bivariate co-clustering analysis quantifies the co-clustering between two differently sized gold nanoparticles tagging two different constituents on the intact PM sheets (36, 37). Similar to the univariate nanoclustering protocol described above, intact apical PM sheets of PSA3 cells co-expressing GFP-LactC2 (probing PS lipids) and an RFP-tagged RAS construct were attached to EM grids and fixed with 4% PFA and 0.1% gluaraldehyde. The PM sheets were incubated with 6 nm gold nanoparticles linked to anti-GFP antibody, blocked with 0.2% bovine serum albumin (BSA) and 0.2% fish skin gelatin, then incubated with 2 nm gold conjugated to anti-RFP antibody. ImageJ was used to assign coordinates to the gold nanoparticle. A bivariate K-function analysis tested the null hypothesis that the two populations of gold particles spatially separate from each other. (Eqs. C-F):

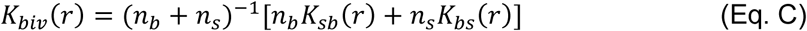

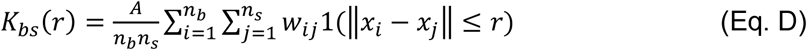

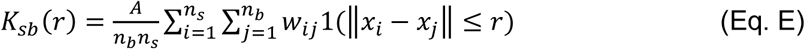

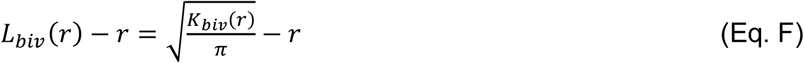

where *K_biv_*(*r*) denotes a bivariate estimator and contains two individual bivariate K-functions: *K_bs_*(*r*) quantifies how the big 6 nm gold particles (*b* = big gold) distribute around each 2 nm small gold particle (*s* = small gold); *K_sb_*(*r*) describes how small gold particles distribute around each big gold particle. The value of n_b_ indicates the number of 6 nm big gold and n_s_ indicates the number of 2nm small gold within a PM area of *A*. Other parameters denote the same definitions as defined in the univariate calculations in Eqs.A and B. *L_biv_*(*r*)-*r* is a linearly transformation of *K_biv_*(*r*), and is normalized against the 95% confidence interval (95% C.I.). An *L_biv_*(*r*)-*r* value of 0 indicates spatial segregation between the two populations of gold particles, whereas an *L_biv_*(*r*)-*r* value above the 95% C.I. of 1 at the corresponding distance of *r* indicates yields statistically significant co-localization at certain distance yields. Area-under-the-curve for each *L_biv_*(*r*)-*r* curves was calculated within a fixed range 10 < *r* < 110 nm, and was termed bivariate *L_biv_*(*r*)-*r* integrated (or LBI):

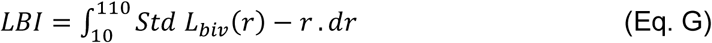

For each condition, > 15 apical PM sheets were imaged, analyzed and pooled, shown as mean of LBI values ± SEM. Statistical significance between conditions was evaluated via comparing against 1000 bootstrap samples as described (36, 37).

### Cell culturing and generation of stable lines

Human and murine pancreatic tumor cell lines, including MOH, PANC1, BxPC3 and iKRAS cells, were maintained in DMEM medium containing 10% fetal bovine serum (FBS). PDAC cell line MiaPaCa-2 was maintained in DMEM medium containing 10% fetal bovine serum (FBS) and 2.5% horse serum (HS). To generate stable cell lines, the pEF6 vector plasmid without/with the cDNA of human LPCAT1 was used to transfect the tumor cells. For each line, 1 μg of plasmid was added to 7 μl of lipofectamine for the transfection. Following 5-hour incubation with the plasmids, cells were washed and changed to DMEM medium containing 10% FBS and 3 μg/mL puromycin antibiotic. Cells were grown in the presence of antibiotics for a week before serial dilution and seeding in 96-well plates with a concentration of < 1 cell per well. Cell colonies were then harvested for Western blotting to verify the expression of LPCAT1.

### Western blotting

Whole-cell lysates of MOH, PANC1 and BxPC3 cells were collected. Following electrophoresis in SDS PAGE gels and transfer, membranes were incubated with primary antibodies against the phosphorylated ERK and Akt, total ERK and Akt, LPCAT1, as well as loading control of actin, overnight. After secondary antibody incubation, membranes were imaged using enhanced chemiluminescence (ECL) solution. Data are shown as mean ± SEM. ImageJ software analysis was used to evaluate expression intensity and identify fold change.

### Proliferation

CyQUANT cell proliferation assay was used to measure number of live cells in microplates. Appropriate number of PDAC cells, such as MOH (1000 cells/well), MiaPaCa-2 (2000 cells/well) and BxPC3 (3000 cells/well), were seeded in 96-well plates. After 96 hours, cells were washed and stained with CyQUANT^®^ GR dye. Following lysis, fluorescence of dye bound to intact nucleic acids was measured using a Tecan plate reader. For each condition, 3 independent experiments were performed, and data were pooled together. Student’s t-test was used to evaluate the statistical significance.

### Colony formation

All human and murine pancreatic cancer cell lines stably expressing V2 or LPCAT1 were seeded in 6-well plates. Specifically, iKRAS (400 cells/well), MOH (250 cells/well) cells were grown for 7 days. MiaPaCa-2 (100 cells/well) were grown for 10 days and BxPC3 (1000 cells/well) were grown for 14 days. Cells were washed twice with PBS, followed with fixation with 4% paraformaldehyde for 15 min. Cell staining was performed with 0.01% crystal violet for 15 min. Colony images were captured using Perkin Elmer X3 multiplate reader. Colony count was performed using ImageJ. For each condition, 3 independent experiments were conducted. Student’s t-test was used to

## Supplemental Figures

**Supplemental Figure 1.** Stable expression of LPCAT1 was verified in MiaPaCa-2 and BHK cells. Whole-cell lysates of BHK and MiaPaCa-2 cells stably expressing V2 empty vector or LPCAT1 were collected for Western blotting. Antibodies against FLAG tag in BHK cells (A) and against LPCAT1 (B) were used and shown.

**Supplemental Figure 2.** LPCAT1 expression significantly elevates the saturated PC and PE levels in MiaPaCa-2 cells. **Whole**-cell lysates of MiaPaCa-2 cells stably expressing V2 or LPCAT1 were collected for lipidomics. Individual species of the saturated PC (A) and PE (B) are shown. (C) Individual PS species in MiaPaCa-2 cells expressing V2 or LPCAT1 are shown. Data are shown as mean ± SEM pooled from 3 independent experiments. Student’s t-test was used to evaluate the statistical significance with * indicating p < 0.05.

**Supplemental Figure 3.** LPCAT1 differentially modulates lipid profiles of MiaPaCa-2 cells. Levels of various lipid types with different numbers of double bonds are shown. Data are shown as mean ± SEM pooled from 3 independent experiments. Student’s t-test was used to evaluate the statistical significance with * indicating p < 0.05.

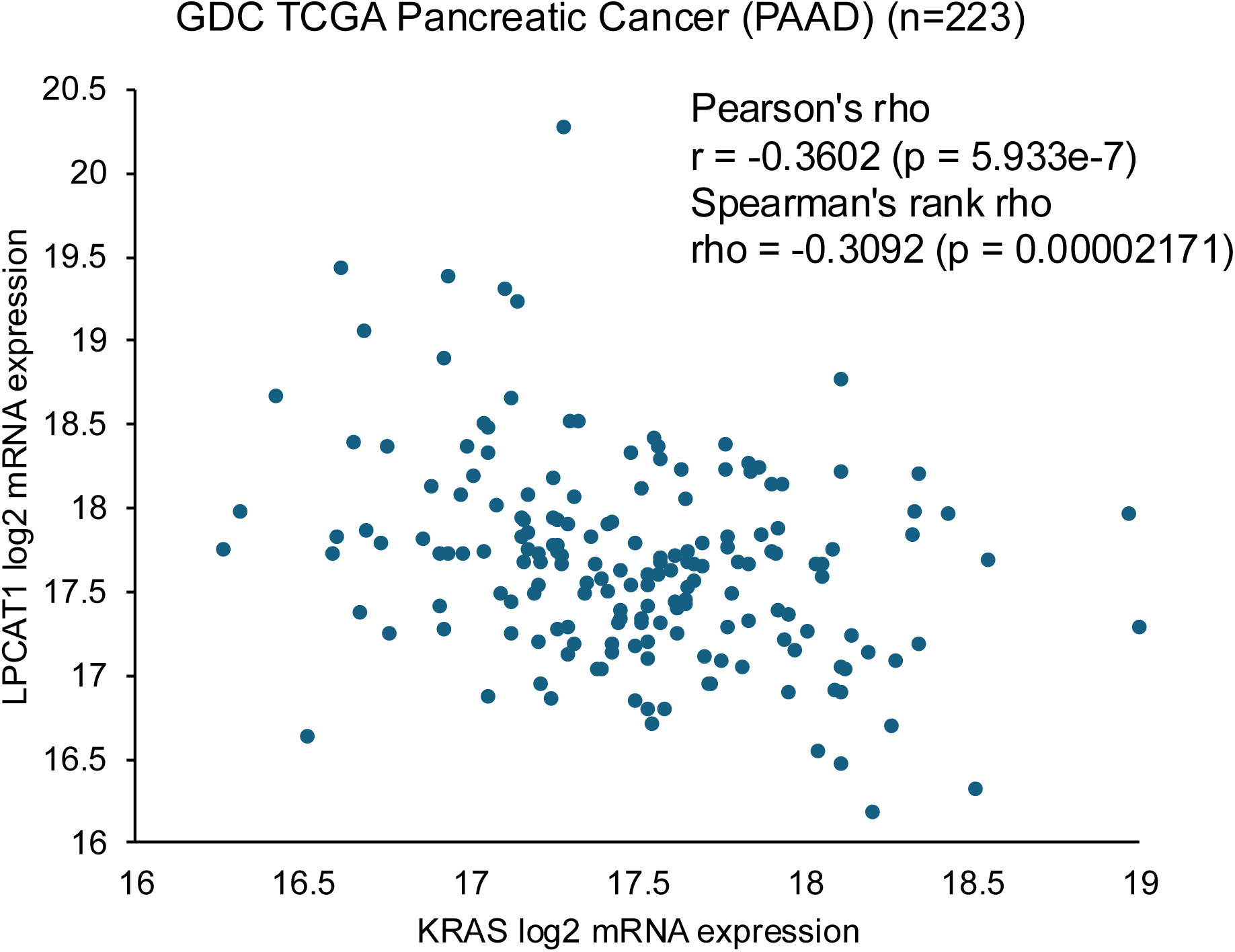

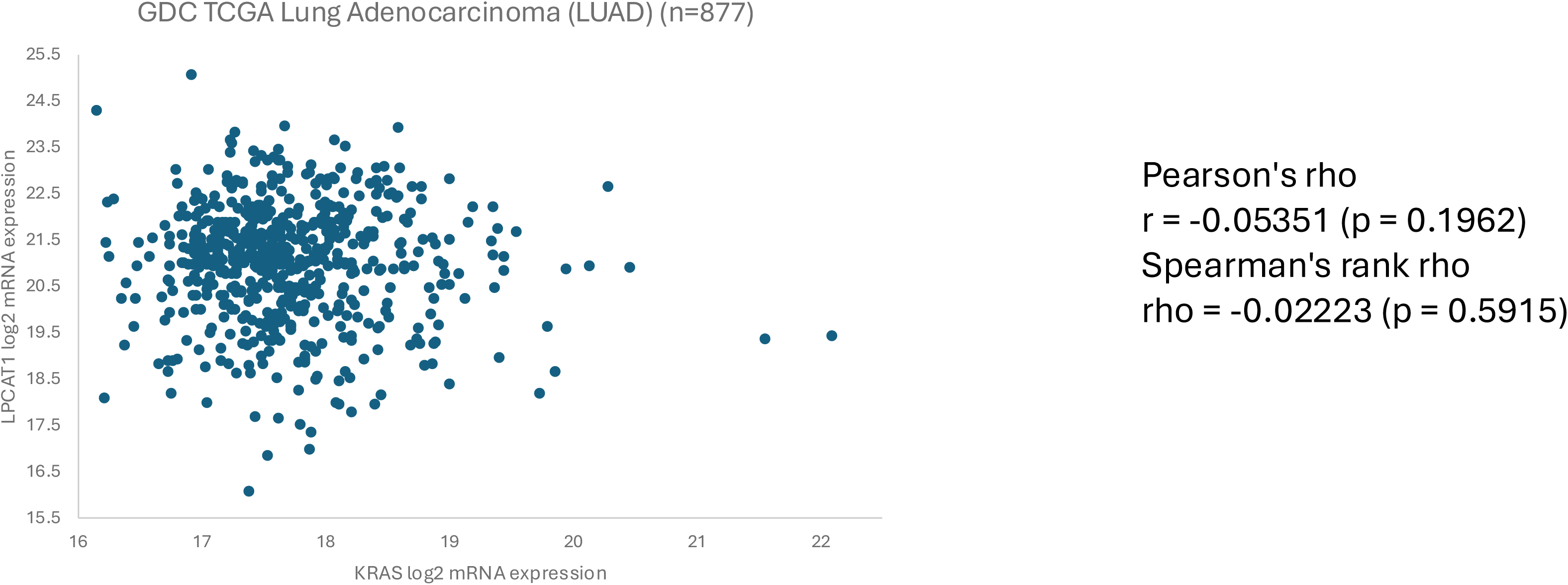

## Bibliography

1. McCormick, F. 2019. Progress in targeting RAS with small molecule drugs. Biochem J 476: 365–374.

2. Cox, A. D., C. J. Der, and M. R. Philips. 2015. Targeting RAS Membrane Association: Back to the Future for Anti-RAS Drug Discovery? Clinical cancer research : an official journal of the American Association for Cancer Research 21: 1819–1827.

3. Cox, A. D., S. W. Fesik, A. C. Kimmelman, J. Luo, and C. J. Der. 2014. Drugging the undruggable RAS: Mission possible? Nature reviews. Drug discovery 13: 828–851.

4. Prior, I. A., F. E. Hood, and J. L. Hartley. 2020. The Frequency of Ras Mutations in Cancer. Cancer Res 80: 2969–2974.

5. Grant, B. J., S. Lukman, H. J. Hocker, J. Sayyah, J. H. Brown, J. A. McCammon, and A. A. Gorfe. 2011. Novel allosteric sites on Ras for lead generation. PLoS One 6: e25711.

6. Ostrem, J. M., U. Peters, M. L. Sos, J. A. Wells, and K. M. Shokat. 2013. K-Ras(G12C) inhibitors allosterically control GTP affinity and effector interactions. Nature 503: 548–551.

7. Arora, N., H. Mu, H. Liang, W. Zhao, and Y. Zhou. 2024. RAS G-domains allosterically contribute to the recognition of lipid headgroups and acyl chains. J Cell Biol 223.

8. Prior, I. A., C. Muncke, R. G. Parton, and J. F. Hancock. 2003. Direct visualization of Ras proteins in spatially distinct cell surface microdomains. J Cell Biol 160: 165–170.

9. Liu, J., N. Arora, and Y. Zhou. 2023. RAS GTPases and Interleaflet Coupling in the Plasma Membrane. Cold Spring Harb Perspect Biol 15.

10. Zhou, Y., and J. F. Hancock. 2023. RAS nanoclusters are cell surface transducers that convert extracellular stimuli to intracellular signalling. FEBS Lett.

11. Prakash, P., D. Litwin, H. Liang, S. Sarkar-Banerjee, D. Dolino, Y. Zhou, J. F. Hancock, V. Jayaraman, and A. A. Gorfe. 2019. Dynamics of Membrane-Bound G12V-KRAS from Simulations and Single-Molecule FRET in Native Nanodiscs. Biophys J 116: 179–183.

12. Prakash, P., Y. Zhou, H. Liang, J. F. Hancock, and A. A. Gorfe. 2016. Oncogenic K-Ras Binds to an Anionic Membrane in Two Distinct Orientations: A Molecular Dynamics Analysis. Biophys J 110: 1125–1138.

13. Neale, C., and A. E. Garcia. 2020. The Plasma Membrane as a Competitive Inhibitor and Positive Allosteric Modulator of KRas4B Signaling. Biophys J 118: 1129–1141.

14. Lee, K. Y., M. Enomoto, T. Gebregiworgis, G. M. C. Gasmi-Seabrook, M. Ikura, and C. B. Marshall. 2021. Oncogenic KRAS G12D mutation promotes dimerization through a second, phosphatidylserine-dependent interface: a model for KRAS oligomerization. Chem Sci 12: 12827–12837.

15. Lee, K. Y., Z. Fang, M. Enomoto, G. Gasmi-Seabrook, L. Zheng, S. Koide, M. Ikura, and C. B. Marshall. 2020. Two Distinct Structures of Membrane-Associated Homodimers of GTP- and GDP-Bound KRAS4B Revealed by Paramagnetic Relaxation Enhancement. Angew Chem Int Ed Engl 59: 11037–11045.

16. Cao, S., S. Chung, S. Kim, Z. Li, D. Manor, and M. Buck. 2019. K-Ras G-domain binding with signaling lipid phosphatidylinositol (4,5)-phosphate (PIP2): membrane association, protein orientation, and function. J Biol Chem 294: 7068–7084.

17. Li, Z. L., P. Prakash, and M. Buck. 2018. A “Tug of War” Maintains a Dynamic Protein-Membrane Complex: Molecular Dynamics Simulations of C-Raf RBD-CRD Bound to K-Ras4B at an Anionic Membrane. ACS Cent Sci 4: 298–305.

18. Lu, H., and J. Marti. 2022. Predicting the conformational variability of oncogenic GTP-bound G12D mutated KRas-4B proteins at zwitterionic model cell membranes. Nanoscale 14: 3148–3158.

19. Lu, H., and J. Marti. 2020. Long-lasting Salt Bridges Provide the Anchoring Mechanism of Oncogenic Kirsten Rat Sarcoma Proteins at Cell Membranes. J Phys Chem Lett 11: 9938–9945.

20. Mazhab-Jafari, M. T., C. B. Marshall, M. J. Smith, G. M. Gasmi-Seabrook, P. B. Stathopulos, F. Inagaki, L. E. Kay, B. G. Neel, and M. Ikura. 2015. Oncogenic and RASopathy-associated K-RAS mutations relieve membrane-dependent occlusion of the effector-binding site. Proc Natl Acad Sci U S A 112: 6625–6630.

21. Simanshu, D. K., M. R. Philips, and J. F. Hancock. 2023. Consensus on the RAS dimerization hypothesis: Strong evidence for lipid-mediated clustering but not for G-domain-mediated interactions. Mol Cell 83: 1210–1215.

22. Harayama, T., M. Eto, H. Shindou, Y. Kita, E. Otsubo, D. Hishikawa, S. Ishii, K. Sakimura, M. Mishina, and T. Shimizu. 2014. Lysophospholipid acyltransferases mediate phosphatidylcholine diversification to achieve the physical properties required in vivo. Cell Metab 20: 295–305.

23. Bi, J., T. A. Ichu, C. Zanca, H. Yang, W. Zhang, Y. Gu, S. Chowdhry, A. Reed, S. Ikegami, K. M. Turner, W. Zhang, G. R. Villa, S. Wu, O. Quehenberger, W. H. Yong, H. I. Kornblum, J. N. Rich, T. F. Cloughesy, W. K. Cavenee, F. B. Furnari, B. F. Cravatt, and P. S. Mischel. 2019. Oncogene Amplification in Growth Factor Signaling Pathways Renders Cancers Dependent on Membrane Lipid Remodeling. Cell Metab 30: 525–538 e528.

24. Akagi, S., N. Kono, H. Ariyama, H. Shindou, T. Shimizu, and H. Arai. 2016. Lysophosphatidylcholine acyltransferase 1 protects against cytotoxicity induced by polyunsaturated fatty acids. FASEB J 30: 2027–2039.

25. Drosten, M., A. Dhawahir, E. Y. Sum, J. Urosevic, C. G. Lechuga, L. M. Esteban, E. Castellano, C. Guerra, E. Santos, and M. Barbacid. 2010. Genetic analysis of Ras signalling pathways in cell proliferation, migration and survival. EMBO J 29: 1091–1104.

26. Ying, H., A. C. Kimmelman, C. A. Lyssiotis, S. Hua, G. C. Chu, E. Fletcher-Sananikone, J. W. Locasale, J. Son, H. Zhang, J. L. Coloff, H. Yan, W. Wang, S. Chen, A. Viale, H. Zheng, J. H. Paik, C. Lim, A. R. Guimaraes, E. S. Martin, J. Chang, A. F. Hezel, S. R. Perry, J. Hu, B. Gan, Y. Xiao, J. M. Asara, R. Weissleder, Y. A. Wang, L. Chin, L. C. Cantley, and R. A. DePinho. 2012. Oncogenic Kras maintains pancreatic tumors through regulation of anabolic glucose metabolism. Cell 149: 656–670.

27. Burge, R. A., and G. A. Hobbs. 2022. Not all RAS mutations are equal: A detailed review of the functional diversity of RAS hot spot mutations. Adv Cancer Res 153: 29–61.

28. Hunter, J. C., A. Manandhar, M. A. Carrasco, D. Gurbani, S. Gondi, and K. D. Westover. 2015. Biochemical and Structural Analysis of Common Cancer-Associated KRAS Mutations. Mol Cancer Res 13: 1325–1335.

29. Roy, S., S. Plowman, B. Rotblat, I. A. Prior, C. Muncke, S. Grainger, R. G. Parton, Y. I. Henis, Y. Kloog, and J. F. Hancock. 2005. Individual palmitoyl residues serve distinct roles in H-ras trafficking, microlocalization, and signaling. Mol Cell Biol 25: 6722–6733.

30. Abankwa, D., A. A. Gorfe, K. Inder, and J. F. Hancock. 2010. Ras membrane orientation and nanodomain localization generate isoform diversity. Proc Natl Acad Sci U S A 107: 1130–1135.

31. Abankwa, D., M. Hanzal-Bayer, N. Ariotti, S. J. Plowman, A. A. Gorfe, R. G. Parton, J. A. McCammon, and J. F. Hancock. 2008. A novel switch region regulates H-ras membrane orientation and signal output. EMBO J 27: 727–735.

32. Cox, A. D., M. M. Hisaka, J. E. Buss, and C. J. Der. 1992. Specific isoprenoid modification is required for function of normal, but not oncogenic, Ras protein. Mol Cell Biol 12: 2606–2615.

33. Hancock, J. F., K. Cadwallader, and C. J. Marshall. 1991. Methylation and proteolysis are essential for efficient membrane binding of prenylated p21K-ras(B). EMBO J 10: 641–646.

34. Baines, A. T., D. Xu, and C. J. Der. 2011. Inhibition of Ras for cancer treatment: the search continues. Future Med Chem 3: 1787–1808.

35. Schultz, D. F., D. D. Billadeau, and S. D. Jois. 2023. EGFR trafficking: effect of dimerization, dynamics, and mutation. Front Oncol 13: 1258371.

36. Zhou, Y., C. O. Wong, K. J. Cho, D. van der Hoeven, H. Liang, D. P. Thakur, J. Luo, M. Babic, K. E. Zinsmaier, M. X. Zhu, H. Hu, K. Venkatachalam, and J. F. Hancock. 2015. SIGNAL TRANSDUCTION. Membrane potential modulates plasma membrane phospholipid dynamics and K-Ras signaling. Science 349: 873–876.

37. Zhou, Y., P. Prakash, H. Liang, K. J. Cho, A. A. Gorfe, and J. F. Hancock. 2017. Lipid-Sorting Specificity Encoded in K-Ras Membrane Anchor Regulates Signal Output. Cell 168: 239–251 e216.

