## Supplementary figures and images for "Nanoclustering and signaling of KRAS G12C and KRAS G12D respond to lipid acyl chain remodeling in an allele-specific manner"

### Supplemental figures

##### Supplemental Figure 1

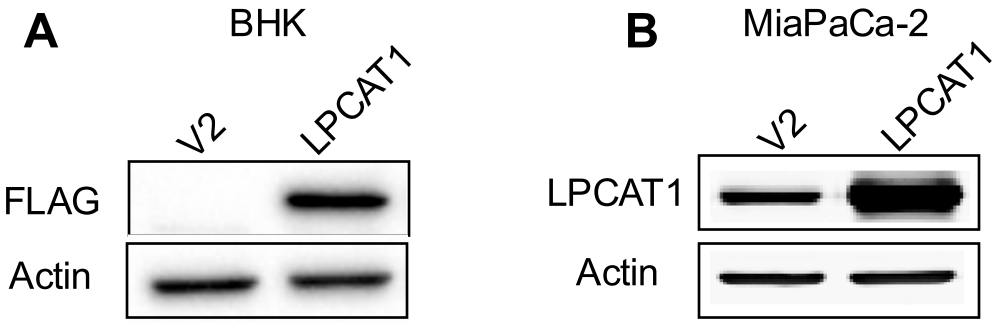

#### Supplemental Figure 2

A

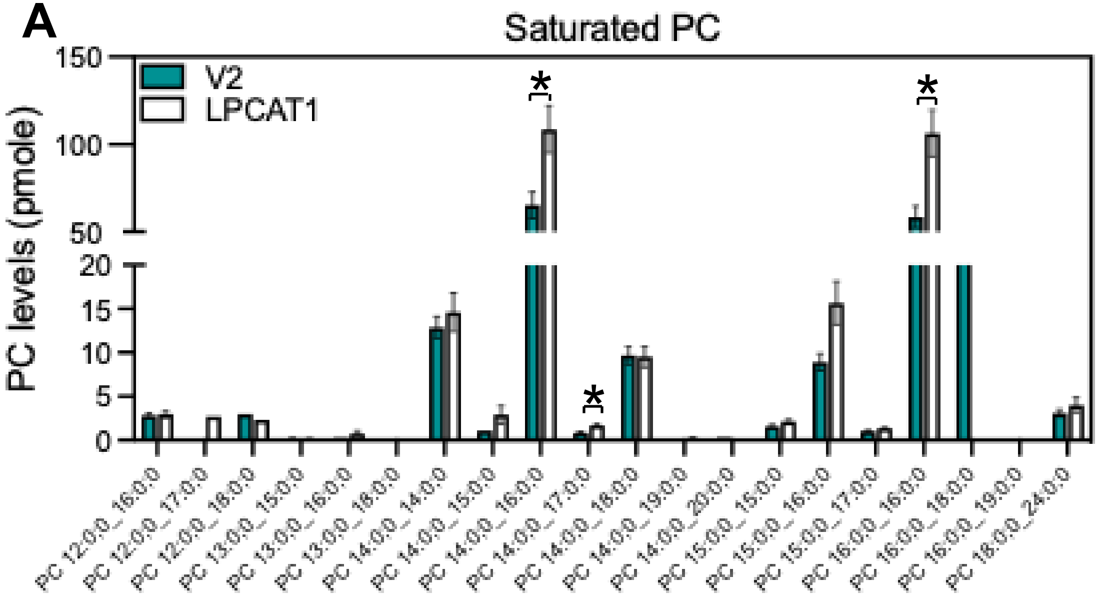


B

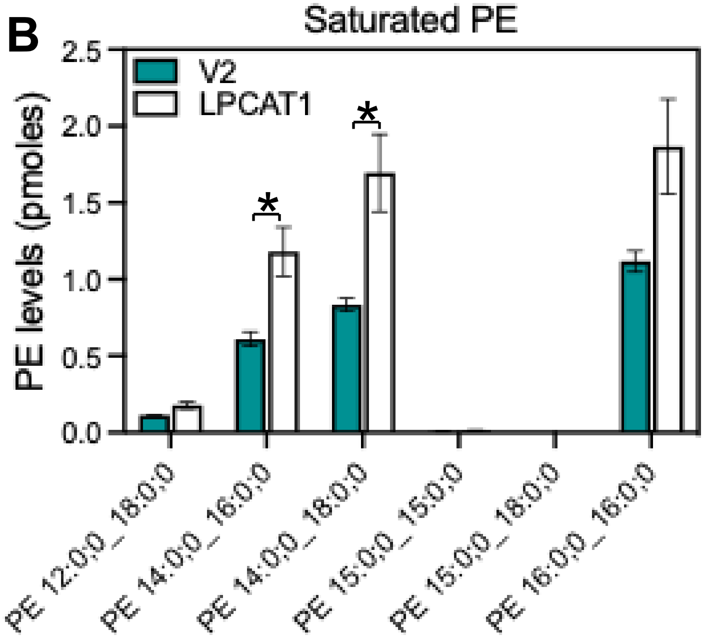

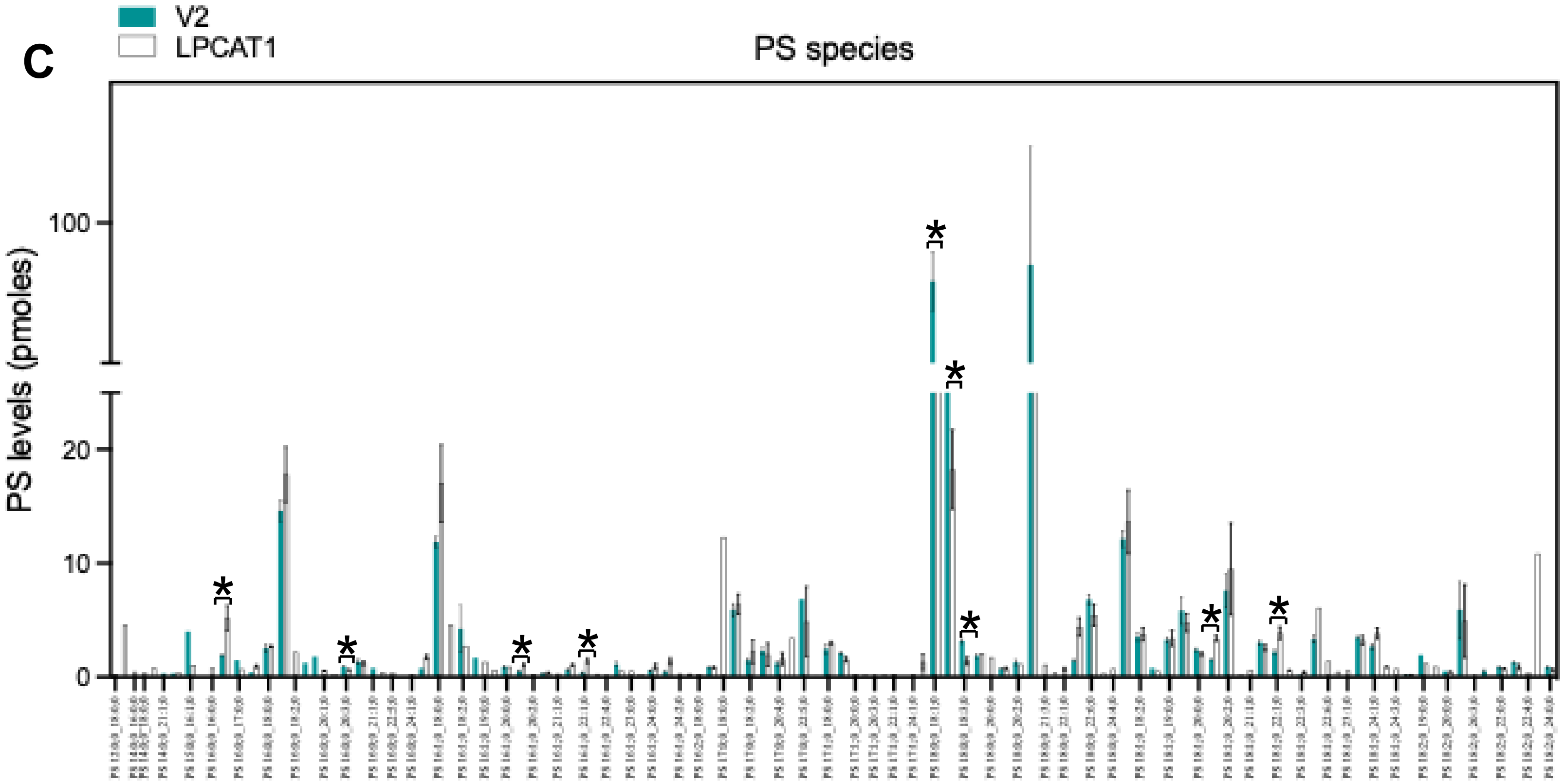

**Supplemental Figure 3**

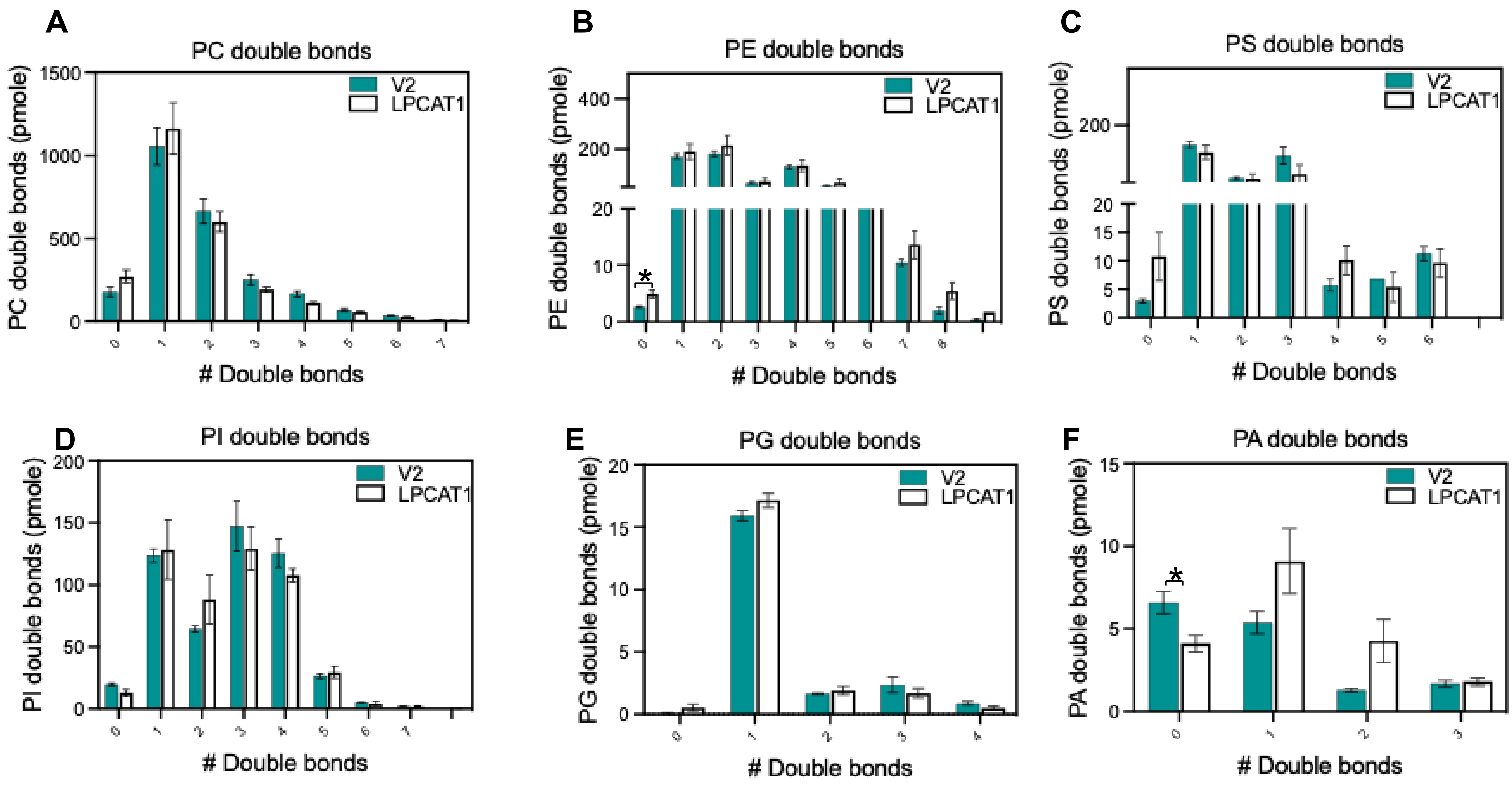

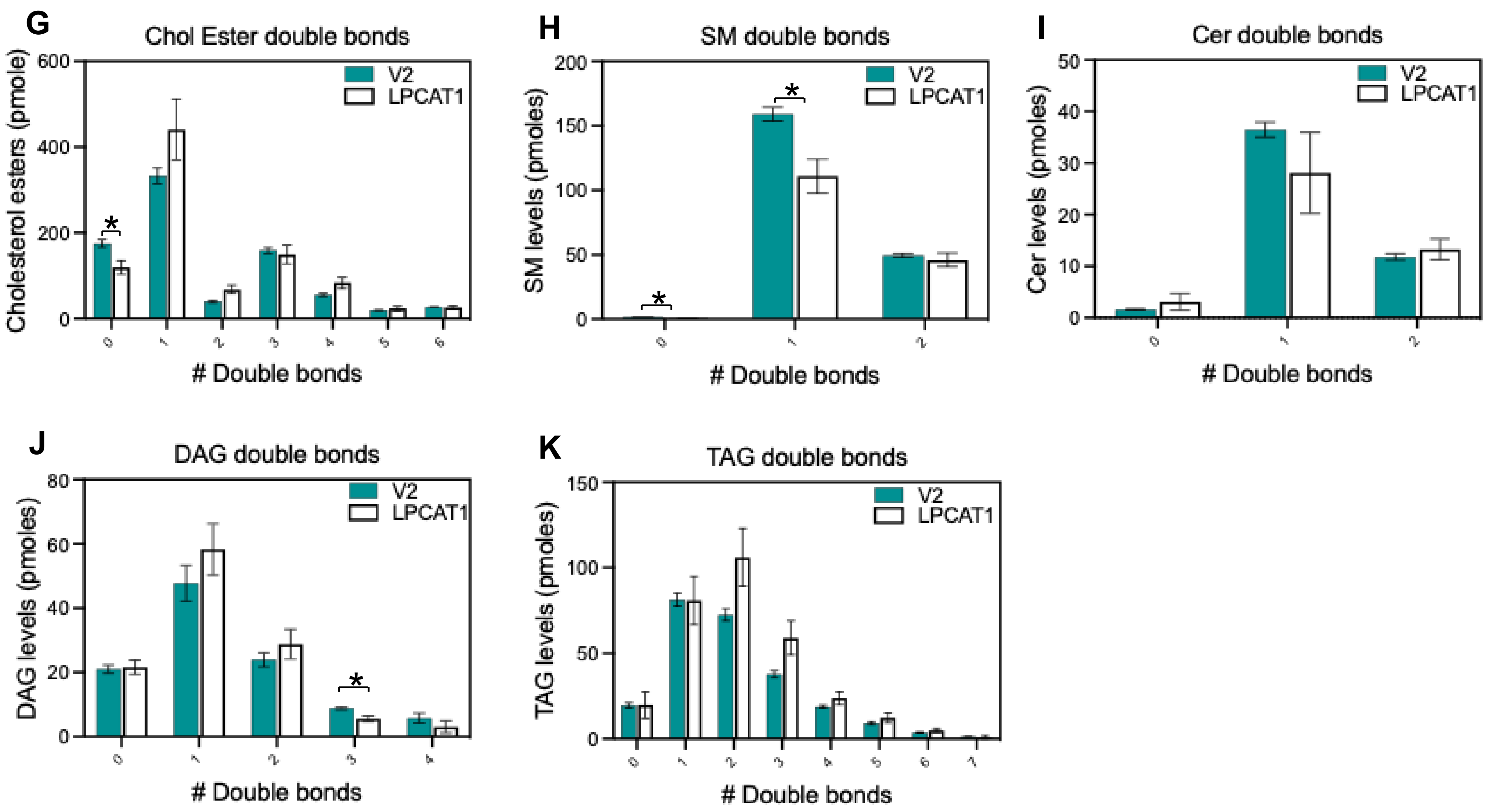
